# A Comprehensive Allele Specific Expression Resource for the Equine Transcriptome

**DOI:** 10.1101/2023.12.31.573798

**Authors:** Harrison D. Heath, Sichong Peng, Tomasz Szmatola, Rebecca R. Bellone, Theodore Kalbfleisch, Jessica L. Petersen, Carrie J. Finno

## Abstract

**Background:** Allele-specific expression (ASE) analysis provides a nuanced view of cis-regulatory mechanisms affecting gene expression.

**Results:** In this work, we introduce and highlight the significance of an equine ASE analysis, containing integrated long- and short-read RNA sequencing data, along with insight from histone modification data, from four healthy Thoroughbreds (2 mares and 2 stallions) across 9 tissues.

**Conclusions:** This valuable publicly accessible resource is poised to facilitate investigations into regulatory variation in equine tissues and foster a deeper understanding of the impact of allelic imbalance in equine health and disease at the molecular level.

## Background

In diploid mammalian cells, autosomal genes are usually equally expressed.^1^ In some cases however, a gene can exhibit expression biased for one allele over the other.^2^ This allele-specific expression (ASE) often results from cis-acting genetic variants on the same chromosome that are in proximity to or within the affected gene. Genetic variants within a gene can affect gene expression, mRNA stability, or mRNA function in different ways, leading to one allele being expressed more than the other. Although not changing the amino acid, synonymous variants, could affect mRNA stability, splicing, or translational efficiency, potentially causing ASE.^3^ Missense variants result in a different amino acid and can affect the function of the RNA, possibly leading to a difference in expression between alleles.^3^ Variants in the 3’ untranslated region (UTR) can influence mRNA stability, localization, and translation, all of which can contribute to ASE. Variants identified in the 5’ prime UTR could influence ASE by affecting the initiation of translation and the stability of mRNA thus altering the amount of protein produced from each allele.^4^ Lastly, variants in splice regions can impact allele expression by altering splicing efficiency, exon skipping, creation or loss of splice sites, or affecting regulatory protein binding, all potentially influencing mRNA function and expression.^4^ ASE may also arise from trans effects arising from genetic influences on the other chromosome of an affected gene or elsewhere in the genome.^5, 6^ Epigenetic factors, such as DNA methylation or chromatin structure, also have the potential to significantly impact gene expression between alleles.^1, 6^ Interestingly, ASE predominantly manifests as tissue-specific phenomena, with loci displaying distinct expression patterns across different tissues. ^2, 5^ Therefore, ASE analysis can provide a way to inspect gene regulation patterns and their impact on biological pathways in specific tissues.

Advances in next-generation sequencing technologies, particularly RNA-sequencing (RNA-Seq), have revolutionized our ability to analyze gene expression and genetic variation. Furthermore, long-read RNA-Seq allows for the straightforward identification of variants that are inherited together as haplotypes.^7, 8^ The identified loci of these haplotypes can then be overlaid with short-read RNA-Seq reads. This integration provides the read counts of nucleotides existing at each locus. These counts can then be used to quantify the expression level of each allele for each gene that contains heterozygous loci. Finally, expression levels of each allele can be compared against one another to identify an allelic imbalance.

In this study, we performed an ASE analysis of the equine transcriptome using a combinatorial RNA sequencing approach. Our aim was to improve the genome-wide functional annotation by providing a comprehensive resource of allelic imbalance detected from linked variants. We provide this analysis to propel future research into important regulatory variants that may impact equine health and disease at the molecular level. This effort is bolstered by the Functional Annotation of the Animal Genome (FAANG) project, a large-scale collaborative effort aimed to identify all functional elements for animals.^9^ The study of ASE in horses is enabled due to the availability of a high-quality reference genome^11^ and long- and short-read sequencing technologies contributing to a more comprehensive transcriptome annotation.^11, 12^ By extending the examination of ASE to the equine species using this method, we can further understand the uses of emerging next generation sequencing technologies and the mechanisms underlying tissue-specific gene regulation in the equine genome.

## Results

Across all samples, we identified 87,174 coding sequence variants. These variants constituted 42,900 haplotypes. After filtering and performing statistical analyses described in *Methods*, we identified 681 ASE events. ASE was found to occur in each tissue analyzed, with the liver and adipose tissues containing the highest frequencies of ASE events. (**Table 1**) Across all samples and tissues in this study, 105 genes exhibited ASE more than once (**Figure 1**).

**Figure 1.**
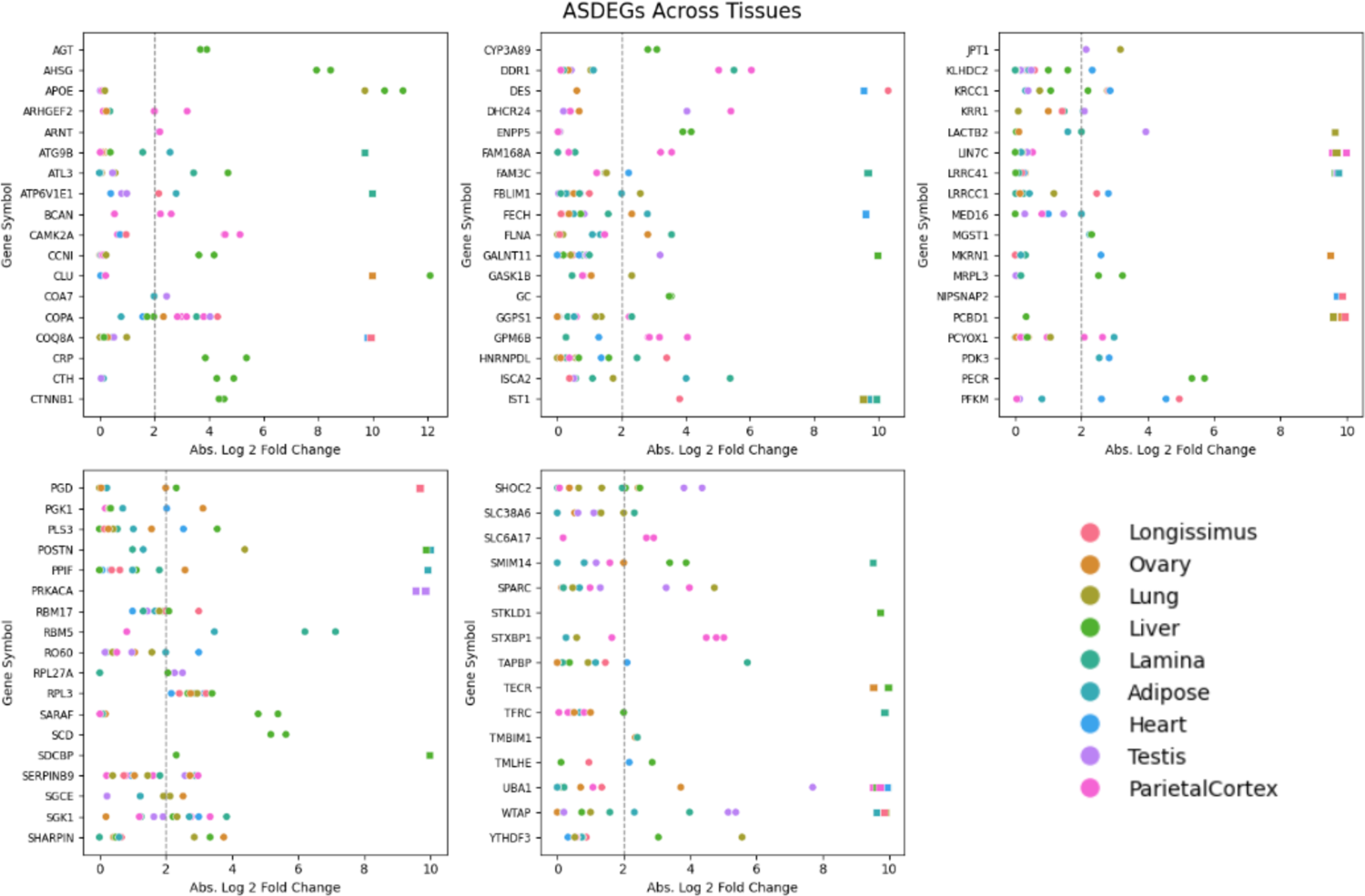
ASDEGs Comparisons Across Tissues. The scatter plots display the absolute log2-fold change in allele expression for identified ASDEGs across various tissues. Each gene featured has at least 2 identified ASE events across all horses and tissues. ASDEGs are graphed in alphabetical order. Each dot (common ASE) or square (mono-allelic expression) represents the expression fold change for a gene in a specific tissue, plotted against the gene symbol on the x-axis and the absolute log2-fold change on the y-axis. In order to visualize genes exhibiting MAE, aeFC was set to a random variable between 9.7 and 10 since an allele expression value compared against zero could not yield a fold change expression level. The dotted line indicates the significance threshold of a 2-fold change. The color coding corresponds to different tissues, as indicated in the legend.

**Table 1.**
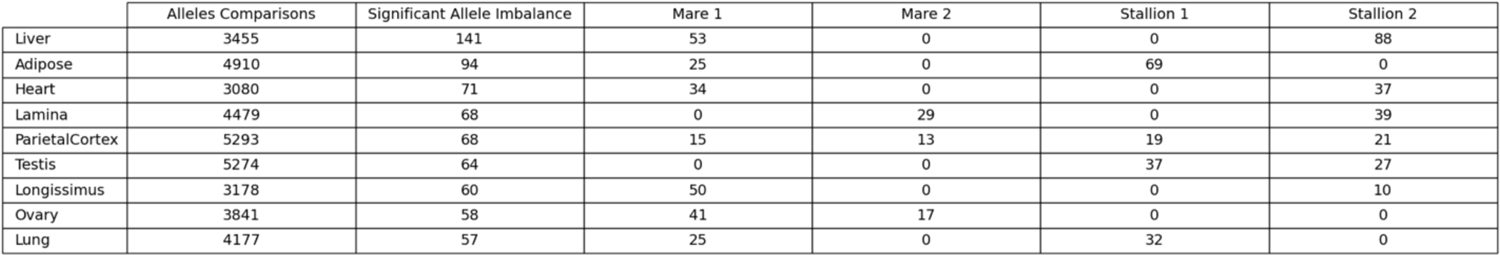
Distribution of Analyzed Genes Across Tissues. An overview of the alleles examined across multiple tissue types delineated by two key metrics. The “Alleles Compared” column enumerates the number of allele comparisons within each specific tissue type (2 alleles for each comparison). The “Significant Allele Imbalance” column identifies the subset of these alleles that exhibited notable expression differences from the expected equilibrium in our study. The other columns represent the number of ASE events from each sample used in the study. Sequencing data selection is described in Methods. The table is organized in ascending order by the number of alleles exhibiting significant allele imbalance.

### Investigating Allele-Specific Variants

Among the 774 variants identified at heterozygous loci within ASE events, 335 were in 3’ untranslated regions, 135 were missense coding variants, 134 were synonymous coding variants, 131 were in 5’ untranslated regions, 23 were splice region variants, 9 were noncoding transcript exon variants, 4 were stop lost variants, and 3 were stop gained variants (**Table 2**). A total of 497 (72.9%) of the identified variants in ASE events fell within histone modified regions. Variants within H3K27ac peaks were found in 377 (55.3%) of the ASE events. H3K4me1 peaks encompassed the variants in 268 (39.4%) of the ASE events and variants within H3K4me3 peaks made up 293 (43.0%) of the events. Lastly, H3K27me3 intersected with 170 (24.9%) of the variants of the ASE events. From the 497 ASE events identified as having SNPs associated with histone modification regions, 369 (74.2%) showed overlap of multiple histone marks. The three most common overlapping histone modification regions with identified variants were H3K4me3 and H3K27ac, H3K27ac and H3K4me1, and H3K27ac with H3K4me1 and H3K4me3 (Figure 2).

**Figure 2.**
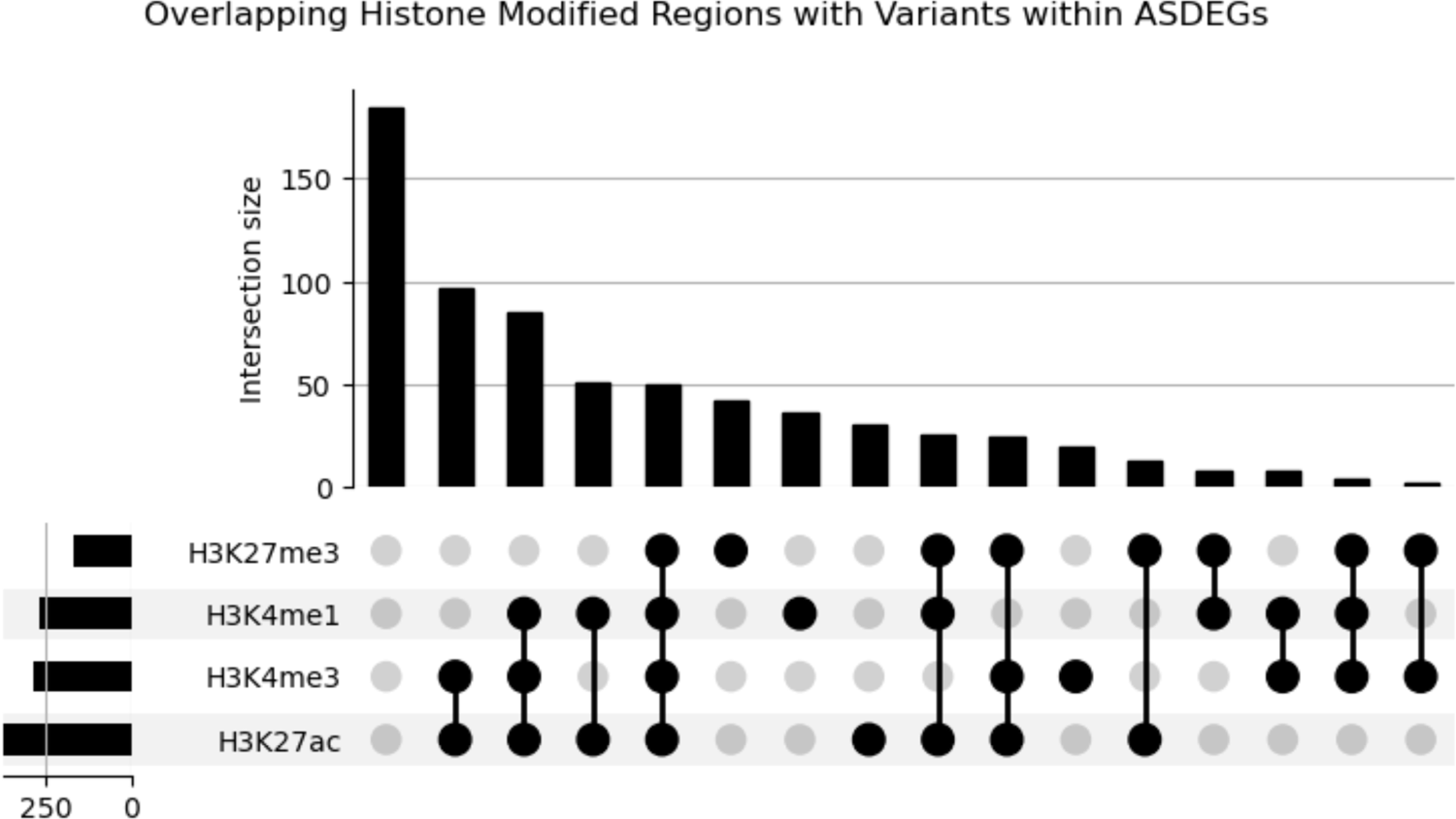
Overlapping Histone Modified Regions with Variants in ASDEGs. Upset plot representing the distribution and overlap of histone modifications across SNP regions associated with allele-specific differentially expressed genes (ASDEGs). Each circle corresponds to a specific histone modification as indicated by the legend (H3K27ac, H3K4me1, H3K27me3, H3K4me3). Circles, and their respective frequency bars, denote the count of SNP regions that exhibit the corresponding histone modifications.

**Table 2.** Variant Types Detected in ASDEGs. Classification of variants caused by heterozygous SNPs within ASE events, as identified by VEP.

|  | Count |
| --- | --- |
| 3 prime UTR variant | 335 |
| missense variant | 135 |
| synonymous variant | 134 |
| 5 prime UTR variant | 131 |
| non coding transcript exon variant | 9 |
| splice region variant & splice polypyrimidine tract variant & intron variant | 6 |
| splice region variant & synonymous variant | 4 |
| missense variant & splice region variant | 4 |
| splice donor region variant & intron variant | 4 |
| stop lost | 4 |
| stop gained | 3 |
| splice region variant & 5 prime UTR variant | 2 |
| splice polypyrimidine tract variant & intron variant | 1 |
| splice region variant & intron variant | 1 |
| missense variant & stop retained variant | 1 |

### Differentially Expressed Gene Enrichment Analysis

We identified 168 KEGG pathways with the potential to be significantly impacted by ASDEGs, including metabolic pathways, endocytosis, and the Ras signaling pathway. In our study, the liver contained the greatest number of pathways significantly impacted by ASE (Figure 3).

**Figure 3.**
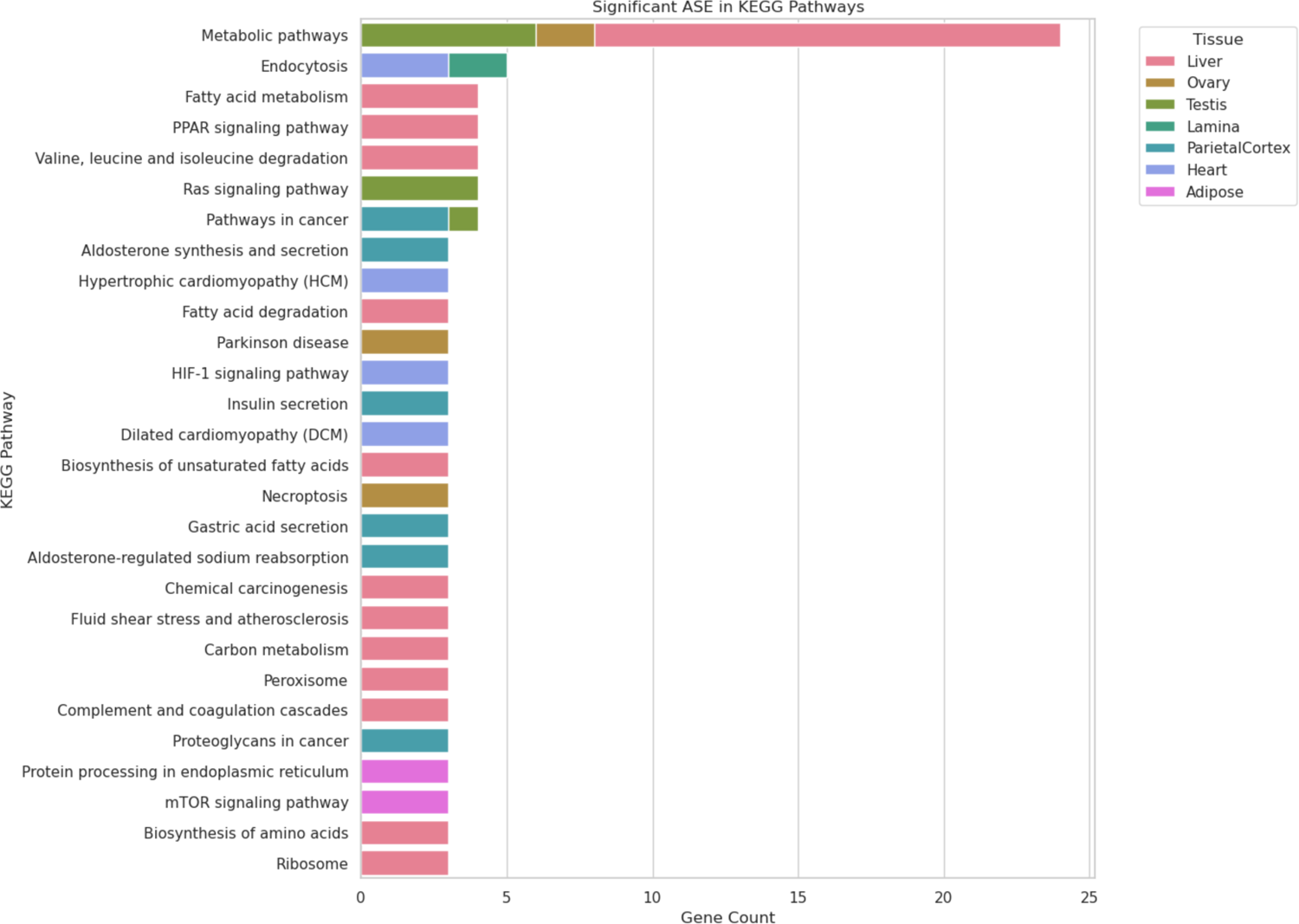
Distribution of Significantly Enriched Pathways Across Different Tissues. The number of ASDEGs present within significantly enriched pathways identified in each tissue type from KOBAS enrichment analysis.

## Discussion

### ASE Analysis

Our ASE analysis began with the identification and assignment of heterozygous variants to specific alleles. The incorporation of long-read RNA-Seq techniques has been instrumental in this aspect. Short-read RNA-Seq’s segmented view of genetic sequences presents challenges in congruent sequence construction.^1^ On the other hand, long-read RNA-Seq’s unfragmented view of the transcriptome facilitates a more robust identification of heterozygous haplotypes.^7, 8^ This methodology surpasses evaluations based solely on expression values of individual heterozygous variants, ensuring a more thorough and accurate assessment.^6–8^ The read counts, provided by short-read RNA-Seq, for each of these positions could then be summed and subsequently averaged by the number of variable loci. Averaging the expression across the variable loci helped to smooth out differences between single position expression values for a more accurately quantified allele expression level.

To reduce the likelihood of including artifacts or minor variations that do not represent true differential expression, we excluded ASE events with variants that were only identified as within intergenic or intronic regions. Given the nature of RNA-Seq data, this filtering ensures that the analysis prioritizes variants that are most pertinent to the corresponding genes. This supplementary data could possibly be used to extend currently annotated coding regions, since this data was generated strictly from high quality RNA-Seq. Additionally, we excluded cases where neither of the allele expression values being compared met or exceeded a threshold of 10 reads. These filtered out events are still available for analysis in supplementary materials.

Then, we focused on determining differential expression between the two alleles of a gene by computing the allelic expression fold change (aeFC). We used this log-transformed fold change to evaluate the relative expression differences because it has been demonstrated to accurately normalize the data from a wide range of expression values.^13^ Relying solely on aeFC, however, may introduce false positives. We tackled this issue by applying a binomial test. The binomial test allows us to confidently identify significant differences between the expression of two alleles. However, given the possibility of type I errors from multiple comparisons, the Benjamini-Hochberg Procedure was applied to the resulting p-values from the binomial test to manage the false discovery rate (FDR). The combined utilization of aeFC and adjusted p-values enabled the development of a dependable allele expression database, providing a platform for further detailed analysis and downstream applications.

### Tissue Regulation via ASE

One of the goals of this study was to identify the role of ASE in tissue specific regulation. We discovered ASDEGs that had ASE in one tissue and equal expression across alleles in other tissues. This suggests that the regulatory mechanisms contributing to these ASDEGs are unique to the particular tissues where they were found.^1, 2^ Such specificity could be due to tissue-specific promoters, enhancers, or other regulatory elements that influence gene expression differently in each tissue type.^1,3^ Apolipoprotein E gene, *APOE*, was among these genes and demonstrated ASE in two out of two liver tissues (one from each sex) used in this study, while having equal allele expression in testis, parietal cortex, and lung tissues. Notably, all ASE events involving *APOE* favored the allele with the same missense variant at locus chr10:15714449. Another gene in this study, *SLC6A17,* is a part of the SLC6 family of transporters. This gene exhibited ASE in 2 out of 4 parietal cortex tissues in this study.

Interestingly, both cases of *SLC6A17* ASE were in parietal cortex tissue of mares, while stallions had a bi-alleleic expression for *SLC6A17* in parietal cortex tissues. Both ASE events involving *SLC6A17* favored the allele with the same 3 prime UTR variant at locus chr5:54600372. Furthermore, neuronal membrane glycoprotein M6-b gene *GPM6B* had ASE in 4 out of 4 samples in parietal cortex tissue used in this study, while having equal allele expression in heart and lamina tissues. All four instances of ASE involving *GPM6B* involved favoring alleles with 3 prime UTR variants (cytosine to thymine) on chrX at positions 10374054, 10374651, and 10375819.

Additionally, we discovered patterns of ASE that were common among multiple tissues suggesting widespread effects on their expression across different tissue types, possibly affecting various biological functions.^1^ Allele specific differentially expressed genes (ASDEGs) demonstrating ASE in more than 4 out of 9 tissues from this study included tubulin gene *UBA1* (9 tissues), L ribosomal protein *RPL3* (8 tissues), serine/threonine protein kinase gene *SGK1* (6 tissues), *WT1*-associating gene *WTAP* (5 tissues), Coat Complex Subunit Alpha gene *COPA* (5 tissues), and lysine-rich coiled-coil protein *KRCC1* (5 tissues).

We also inspected instances presenting signs of monoallelic expression (MAE), a phenomenon where only one of the two alleles of a gene is actively transcribed.^14^ MAE was identified by the event where one allele had an expression value of 0 and the other had expression >= 5. This decision was made to account for the chance of sequencing or sampling error in short read RNA-Seq without introducing too many false positives. Out of our identified ASE events, 119 (17.4%) showed statistical evidence of mono-allelic expression (MAE). Tubulin gene *UBA1* exhibited MAE in 8 out of 9 tissues used in our study. Notably, each of these events favored the allele with missense variants (cytosine to thymine) on chrX positions 39703029 and 39703030. Additionally, all of these MAE events involving *UBA1* were in tissues from stallions. In this study, *UBA1* showed equal expression across alleles in 8 out of 9 tissues for mares.

The allele expression data frame resulting from this analysis enables researchers to look up genes of interest and is publicly available in the data and materials section. Additionally, this main database was broken up into tissue specific tracts allowing for tissue-specific allele expression analysis.

### Histone Modifications

ChIP-seq data was integrated to provide a broader evaluation of allele-specific expression in the context of annotated histone modifications. This leads to an improved perspective on the epigenetic acting influences in allele-specific gene expression, illuminating the complex interplay between genetic variations and epigenetic regulation. Variants within H3K27ac peaks, typically present at active transcription start sites (TSS), highlight the potential relationship between coding changes within TSS and allelic expression.^15^ Additionally, intersection of ASE events with H3K4me1 peaks, often associated with enhancer regions, suggests that variants within these regions could lead to enhanced expression relative to the other allele.^15^ Variants were also identified within H3K4me3 peaks, which are commonly identified near promoters of actively transcribed genes. This suggests that a particular promoter sequence may be favored for transcription.^15^ Similarly the repressive mark, H3K27me3, further emphasizes the possible interplay between allele imbalance and the epigenetic landscape.^15^ The co-localization of ASE with active marks such as H3K27ac and H3K4me1, often found at transcription start sites and enhancer regions, respectively, suggests that these epigenetically active domains may predispose certain alleles for increased expression by facilitating a more accessible chromatin state.^20^ Simultaneously, the intersection with H3K4me3, associated with promoters of actively transcribed genes, and H3K27me3, a mark of transcriptional repression, indicates a nuanced regulatory landscape where alleles may be differentially expressed due to the combinatorial effects of epigenetic modifications.^15^ These overlapping epigenetic regions may serve as hotspots for ASE, where the orchestration of gene activation and silencing is fine-tuned by histone marks and coding variants. However, it is important to note that not all impactful epigenetic markers can be found in transcribed regions.

### Gene Enrichment Analysis

This ASE analysis also provided insights into cellular processes that are putatively modulated by allele expression imbalances. This comprehensive view, which combines data from multiple samples sourced from the same tissue type, can reveal how ASE might broadly affect biological functions.^16^ The resulting database of pathways predicted to be altered within each tissue is a valuable resource for further examination of variants contributing to disease or functional traits.

### Limitations & Future Direction

In this study, we introduce a resource for the equine community designed to advance future gene regulation investigations. Additionally, we presented methodologies for identifying variants potentially responsible for regulation of gene expression, identifying pathways possibly altered by these regulatory events, and characterizing ASE patterns on a tissue-wide scale. However, there are a few limitations to this approach. First, our sample size was relatively small, and we did not analyze all tissues, which may affect the generalizability of our findings. Additionally, we did not experimentally validate candidate ASE genes. It is also important to note that this combinatorial RNA-Seq method simply will not detect all instances of ASE.^17^ Our approach of using heterozygous gene expression markers reduces the total number of ASE events we could identify in the study. as some locations in the genome may be homozygous yet still exhibit ASE due to epigenetic factors alone.^18^ Studies utilizing both RNA-Seq and whole genome sequencing have estimated the percent of genes exhibiting ASE from 1% to 20%, depending on the strength of statistical filtering.^1, 18^ In this study, we used high filtering standards to declare a significant allele imbalance, and identified ASE in 1.59% of genes analyzed. Filtering in this study is depicted in **Supplementary** Figure 1. While we may not be able to detect all ASE events, using this method provides a helpful way to inspect linked variants and their putatively high impact on gene expression. Future investigations will extend this approach to a larger set of samples and tissues. Additionally, experimental validation into these candidate genes and incorporation of other types of next-generation sequencing data will further reveal what is driving ASE. This comprehensive integration will lay the foundation for a deeper understanding of the intricate relationship between imbalances in allele expression and their consequential impacts on the equine genome.

## Conclusion

This study introduces the first multi-tissue analysis of ASE specific to the equine genome, spanning 4 individual horses and 9 tissues. Additionally, it provides downstream analysis techniques that could be used to identify important regulatory variants, significantly impacted pathways, and tissue-specific function from the generated data. We have demonstrated the potential that this method provides to enrich our understanding of the intricacies of equine genetic regulation contributing to function and disease.

## Methods

### Generation of Data

Data from eight tissues from prior FAANG analyses were selected for ASE analyses: lamina, liver, left lung, left ventricle of the heart, longissimus muscle, skin, parietal cortex, testes, and ovary from 4 healthy Thoroughbreds (2 mares and 2 stallions).^19, 20^ This selection aimed to encompass a broad spectrum of biological functions and, consequently, varied gene expression.

The RNA isolation for Iso-seq was performed separately from the same tissues as the RNA utilized for mRNA-Seq, using an identical protocol.^12^ For Iso-seq, we selected the highest quality RNA per sex for each tissue, except for parietal cortex. Since parietal cortex was the pilot tissue, long-read RNA sequencing was done and analyzed for all four horses. All selected RNA samples had integrity (RIN) values greater than or equal to 7. Selected tissues were processed for Iso-seq in a single batch. The cDNA libraries were developed and sequenced at the UC Berkeley QB3 Genomics core facility. Two randomly selected libraries were combined and then sequenced together on a single SMRT cell of the PacBio Sequel II system, as previously described.^11, 15^ ChIP-seq data for the histone modifications evaluated in this study were sourced and analyzed from prior publications that examined the same eight tissues.^11, 21^

### Identifying Haplotypes

The overall workflow for identifying ASE events is outlined in **Supplementary** Figure 2. Key to our approach was the use of Iso-seq data to extrapolate haplotypes for each horse individually, utilizing the isophase Cupcake software v29.0.^22^ The software’s function was to identify variations between otherwise identical full-length reads, thereby pinpointing specific positions of heterozygous SNPs. Notably, the software will only declare haplotypes if it reaches a read depth of 40x, ensuring biologically accurate haplotypes.^22^

### Integration of Short-Read RNA Library

To prepare short-read RNA-Seq data for analysis, sequences were trimmed to remove adapters low-quality read, and PCR duplicates using trim-galore v0.6.10^23^ and Cutadapt v4.7.^24^ Read qualities were inspected using fastQC v0.11.7^25^ and multiQC v1.16^26^, and filtered for reads >= 50bp and quality >= 30. From the identified Heterozygous SNP loci, the varying nucleotides expressed at these positions were generated using SAMtools mpileup from SAMtools 1.18.^27^ We ensured that both the short-read and long-read RNA-Seq analyses were based on the same tissues and samples.

### Quantify Expression for Heterozygous Loci

After overlaying reads found at the heterozygous loci, we quantified the expressed nucleotides at these positions. This was accomplished using a custom Python tool.^28^ The fundamental principle behind this tool is to tally the occurrences of each nucleotide variant at a given locus. For example, if a particular position has both ‘A’ and ‘G’ variants, the tool calculates the frequencies of ‘A’ and ‘G’ in the reads. This process is repeated across all identified heterozygous sites in the genome.

### Quantify Allele Expression Per Haplotype

The next step in our analysis involved aggregating individual nucleotide counts to quantify allele expression. When certain variants within a haplotype appear more frequently in the reads compared to its counterpart, it is evidence that this particular allele is being expressed at a higher level. To carry this out, we first aligned the expression values with the sequence of DNA variants that were identified as haplotypes based on the long-read Iso-seq data. Read counts per locus were then summed according to their respective haplotypes. Then, we divided the combined read counts by the number of variable loci in the haplotype giving what we referred to as allele expression values. The distribution of read counts across all samples is provided in supplementary materials (**Supplementary** Figure 3).

### Identify Significant Allele Specific Expression

After deriving allele expression, we then pinpointed significant ASE using filtering and statistical techniques outlined below. First, we excluded cases where neither of the allele expression values being compared met or exceeded a threshold of 10. We then assessed ASE by examining the differential expression across haplotypes and subsequently applied a log transformation to calculate allele expression fold change (aeFC). The equation for aeFC is shown below:

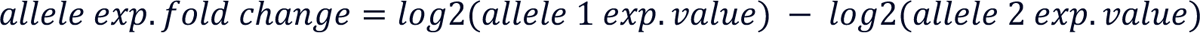

The distribution of aeFC across all samples is provided in supplementary materials (Supplementary Figure 4).

Operating under the null hypothesis that ASE values for a specific gene should be roughly equivalent between alleles, we calculated p-values using a binomial test, assessing the disparities between each gene’s allele expression value. Considering the count-based nature of our data, we employed the Benjamini-Hochberg procedure to manage the expected false discovery rate, yielding adjusted p-values.

For the final stage of significant ASE identification, we established stringent criteria to distinguish instances of significant allele expression imbalance: the aeFC between two alleles had to be at least an absolute value of 2, and the adjusted p-value determined from AE difference needed to be ≤ .05.

To delineate MAE in our dataset, we established a stringent threshold for allele expression. Specifically, we characterized MAE by the criterion where one of two alleles’ read counts from short-read RNA-Seq was 0, and the other was greater than or equal to 5.

### Incorporating Histone Modification Data

ChIP-seq data was integrated by delineating a region extending from the initial to the last heterozygous position in our haplotype transcripts. Leveraging this defined region, we executed computational assessments to ascertain any overlaps with key chromatin peaks, specifically H3K27me3, H3K27ac, H3K4me1, and H3K4me3, on a tissue and sample basis.

### Identify Genes with ASE

We then identified the genes shown to exhibit ASE. To do so, we employed Ensembl’s Variant Effect Predictor (VEP) v10^28^ on the delineated heterozygous loci that made up each haplotype.

VEP’s online tool uses the latest gene transfer format files to identify genes based on a given locus.^28^ VEP may provide multiple annotations for a single variant, therefore, variants predicted as intronic or intergenic were filtered out to only annotate variants strictly from RNA-Seq. If a variant’s annotation was predicted solely as intergenic or intronic, the data was not included in the downstream analyses.

### Allele Specific Differentially Expressed Gene Enrichment Analysis

In our investigation of allele-specific expression (ASE) and its impact on biological pathways, we performed gene enrichment analysis (GEA) using the KOBAS^29^ (KEGG Orthology-Based Annotation System). Our method involved a deliberate combination of allele-specific differentially expressed genes (ASDEGs) from various samples, but importantly, this was done on a tissue-specific basis. By inputting these consolidated ASDEG lists into KOBAS for each tissue type, accessed via http://kobas.cbi.pku.edu.cn/genelist/, we could identify pathways likely to be influenced by ASE across samples. P-values were supplied by KOBAS for each impacted pathway, and significant pathways were identified as having a p-value less than 0.05. Dataframe management and statistical analyses were performed using scipy^30^, numpy^31^, and pandas^32^. Data visualization was achieved using matplotlib^33^ and seaborn^34^.

## Abbreviations

aeFC: allelic expression fold change

ASE: allele specific expression

ASDEG: allele specific differentially expressed gene

RNA-Seq: RNA-sequencing

SNP: single nucleotide polymorphism

GEA: gene enrichment analysis

MAE: mono-allelic expression

## Declarations

### Ethics approval and consent to participate

All protocols were approved by the University of California Davis Institutional Animal Care and Use Committee (Protocol #19037).

### Consent for publication

Not applicable.

### Availability of data and materials

The master and tissue separated allele expression data frames generated and analyzed in this study are available as **Supplementary Files 3-10**.

The short-read RNA sequencing analyzed in this study is available in the ENA and SRA repositories under the accession number PRJEB26787 (female tissues - https://www.ebi.ac.uk/ena/browser/view/PRJEB26787) and PRJEB53382 (male tissues - https://www.ebi.ac.uk/ena/browser/view/PRJEB53382).

The Iso-seq data analyzed in this study is available in the ENA and SRA repositories under the accession number PRJEB53020. (https://www.ebi.ac.uk/ena/browser/view/PRJEB53020)

Histone ChIP-seq analyzed in this study is available via Kingsley et al. (https://doi.org/10.3390/genes11010003) and Barber et al (thesis; https://digitalcommons.unl.edu/animalscidiss/233/).

## Authors’ Contributions

Substantial contributions to the conception & design of the work, Harrison Heath, Sichong Peng, Carrie J. Finno; the acquisition, analysis, and interpretation of data, Harrison Heath, Sichong Peng, Carrie J. Finno, Tomasz Szmatola; drafting manuscript, Harrison Heath, Sichong Peng, Carrie J. Finno; manuscript review and revisions, Rebecca Bellone, Theodore Kalbfleisch, Jessica L. Petersen, Sichong Peng, Carrie J. Finno. All authors have approved the submitted version and any substantially modified version that involves the author’s contribution to the study, and agreed both to be personally accountable for the author’s own contributions and to ensure that questions related to the accuracy or integrity of any part of the work, even ones in which the author was not personally involved, are appropriately investigated, resolved, and the resolution documented in the literature.

## Acknowledgements

None.

## Supplementary Files

Supplementary Figure 1. Sankey Diagram representing the filtering process used in this study.

Supplementary Figure 2. Depiction of overall pipeline for ASE identification.

Supplementary Figure 3. Distribution of read counts for heterozygous loci used in this study.

Supplementary Figure 4. Bar plot showing the distribution of allele expression differences analyzed in this study. The red dotted line represents the cut off for significant ASE.

Supplementary Table 1. Filtered out Allele expression values.

Supplementary Table 2. Results from VEP.

Supplementary Tables 3-10. The master and tissue separated allele expression data frames generated and analyzed in this study.

### Codes/Scripts

Computer codes/scripts used in this analysis are publicly available on github at: https://github.com/hdheath/ASE_equine_transcriptome/blob/main/README.md

